# Biotic interactions and environmental filtering both determine earthworm alpha and beta diversity in tropical rainforests

**DOI:** 10.1101/2024.10.12.617983

**Authors:** Arnaud Goulpeau, Mickaël Hedde, Pierre Ganault, Emmanuel Lapied, Marie-Eugénie Maggia, Eric Marcon, Thibaud Decaëns

**Affiliations:** CEFE, Univ Montpellier, CNRS, EPHE, IRD, Montpellier, France; Eco&Sols, INRAE, IRD, CIRAD, Institut Agro, Montpellier, France; ECODIV, Université de Rouen, INRAe, Rouen, France; Taxonomia Biodiversity Fund, 7 rue Beccaria, 72012 Paris, France; Department of Integrative Biology, University of Guelph, N1G 2W1 Guelph, Canada; AgroParisTech, UMR AMAP, CIRAD, CNRS, INRAE, IRD, Univ Montpellier, Montpellier, France

**Keywords:** Alpha diversity, beta diversity, community ecology, Oligochaeta, tropical rainforest

## Abstract

1. Understanding the relative importance of biotic interactions, multiple environmental drivers, and neutral processes in shaping community diversity and composition is a central question for both theoretical and applied ecology.
2. We analysed a dataset describing 125 earthworm communities sampled in 10 localities in French Guiana. DNA barcodes were used to delimit operational taxonomic units (OTUs) that we considered as species surrogates to avoid the taxonomic deficit and calculate community-scale species richness and pair-wise Sørensen beta-diversity. We used log-ratio and generalised linear models to highlight the effects of biotic interactions and environment as drivers of alpha diversity, and generalised dissimilarity models to figure out the relative contribution of space and environment to beta-diversity at different spatial extents.
3. Community-scale alpha diversity was mainly explained by habitat filtering (soil texture) and interspecific competition that limit the number of locally co-existing species.
4. Beta diversity between pairs of communities was mainly explained by distance when comparing communities in similar habitats, by topography and available soil phosphorus when comparing communities in different habitats, and by distance, elevation and climate when comparing all possible pairs of communities.
5. While community composition is determined locally by neutral processes and environmental filtering, biogeographic processes linked to dispersal limitation and adaptation to local environment are the most influential on a regional scale. This highlights the complex interplay of dispersal limitation, biotic interactions and environmental filtering during the process of community assembly.

## Introduction

Understanding how and why community diversity and composition vary over space is a question of fundamental and practical interest to a wide range of researchers in ecology, biogeography, evolutionary biology and biodiversity conservation (Dambros et al., 2020; Seymour et al., 2024). An essential tool for describing these spatial biodiversity patterns and understanding their underlying processes is to break down regional species diversity into its alpha and beta components (Baselga, 2010). While alpha diversity is generally influenced by local factors such as resource availability and biotic interactions (Allbee et al., 2023; Parmentier et al., 2007), beta diversity is assumed to reflect species dispersal processes and the effects of large scale environmental gradients and biogeographical barriers (Dambros et al., 2020; Graco-Roza et al., 2022). The comparative study of alpha and beta diversity therefore provides a more comprehensive picture of biodiversity, enabling a better understanding of the effects of global change on ecological communities (Jones et al., 2022; Kessler et al., 2009). This is an essential step towards designing conservation strategies that support both local diversity and the regional resilience of ecosystems (Gardner et al., 2010; Socolar et al., 2016).

The body of literature on biodiversity spatial patterns has focused mainly on a limited number of taxa, essentially plants, vertebrates and a few groups of insects. Soil organisms, which account for up to 59% of global biodiversity and are recognised for their essential role in the functioning of terrestrial ecosystems, have paradoxically been overlooked (Anthony et al., 2023; Decaëns, 2010; Decaëns et al., 2006). Earthworms may seem to be an exception to this situation, since they have been the subject of numerous studies comparing other soil taxa, but these are largely biased towards local studies, in agricultural and essentially temperate environments. Phillips et al. (2019) studied the global distribution of local earthworm richness, showing that it is essentially driven at this scale by climate, more specifically rainfall, and secondly by elevation. Ganault et al. (2021) further suggested that macrodetritivores, including earthworms, may respond to tree composition and the resulting input of dead plant products such as leaf and root litter and rhizodeposition. Phillips et al. (2019) have also shown that local species richness surprisingly peaks at temperate latitudes, displaying a pattern opposite to that classically observed for aboveground organisms. However, they also show that beta diversity is higher at lower latitudes, contributing to maintaining high regional diversity in the tropics.

This trend towards low alpha diversity and high beta diversity explaining outstanding regional diversity is supported by more detailed studies in the tropical rainforests of French Guiana (Decaëns et al., 2016; Goulpeau, Hedde, Ganault, et al., 2024; Maggia et al., 2021). Limited community-scale richness observed in these studies has been hypothesized to result from low diversity of resource types in soil and to the ecological plasticity of most earthworm species, which contribute to a rapid saturation of ecological space during community assembly (Decaëns et al., 2016). Beta diversity seems to be explained by a spatial effect resulting from limited dispersal, and by habitat features, including vegetation composition and soil properties (Decaëns et al., 2016; Maggia et al., 2021; Vleminckx et al., 2019). However, these studies were not designed to formally separate local from regional effects, and we still need a more comprehensive analysis of the driving factors of both species richness and compositional dissimilarity through a hierarchy of clearly identified spatial scales. This would provide new insights into the assembly rules of earthworm communities, while shedding light on the factors responsible for the extraordinary diversity observed in French Guiana’s diversity hotspot.

In this study, we explored a substantial dataset accumulated following several sampling expeditions to tropical rainforests of French Guiana (Goulpeau, Hedde, Maggia, et al., 2024). We used this dataset to identify the contribution of local environment and biotic interactions as drivers of community-scale alpha diversity, and the relative importance of environmental dissimilarity and geographical distance as drivers of beta diversity among pairs of communities. This was done by considering a hierarchy of nested spatial extents. We hypothesised: 1) that community-scale richness is explained by community saturation, which limits the number of locally coexisting species, and by local environmental conditions, such as climate and soil properties; 2) that compositional dissimilarity between communities is essentially explained by environmental heterogeneity at a local scale, and by geographical distance and large scale environmental gradient at a regional scale.

## Material and method

### Study sites

We sampled earthworm communities between 2011 and 2019 in ten localities of French Guiana (Fig. 1; SI Table S1). Climate in the study region is humid tropical, with an average annual temperature of 26 °C and average annual rainfall ranging from 2,000 to 4,000 mm (Barret, 2001). The elevation is generally less than 200 m above sea level, and the region is criss-crossed by a dense network of rivers and dotted with isolated hills and inselbergs that can peak at around 800 m (Barret, 2001). Soils are mainly Ferralsols in well-drained areas, sometimes superficially hardened by laterite crusts, Lithosols on granitic outcrops (inselbergs and other landforms), Acrisols in lowland areas, and Podzols in sandy coastal regions (Ferry et al., 2003).

**Figure 1.**
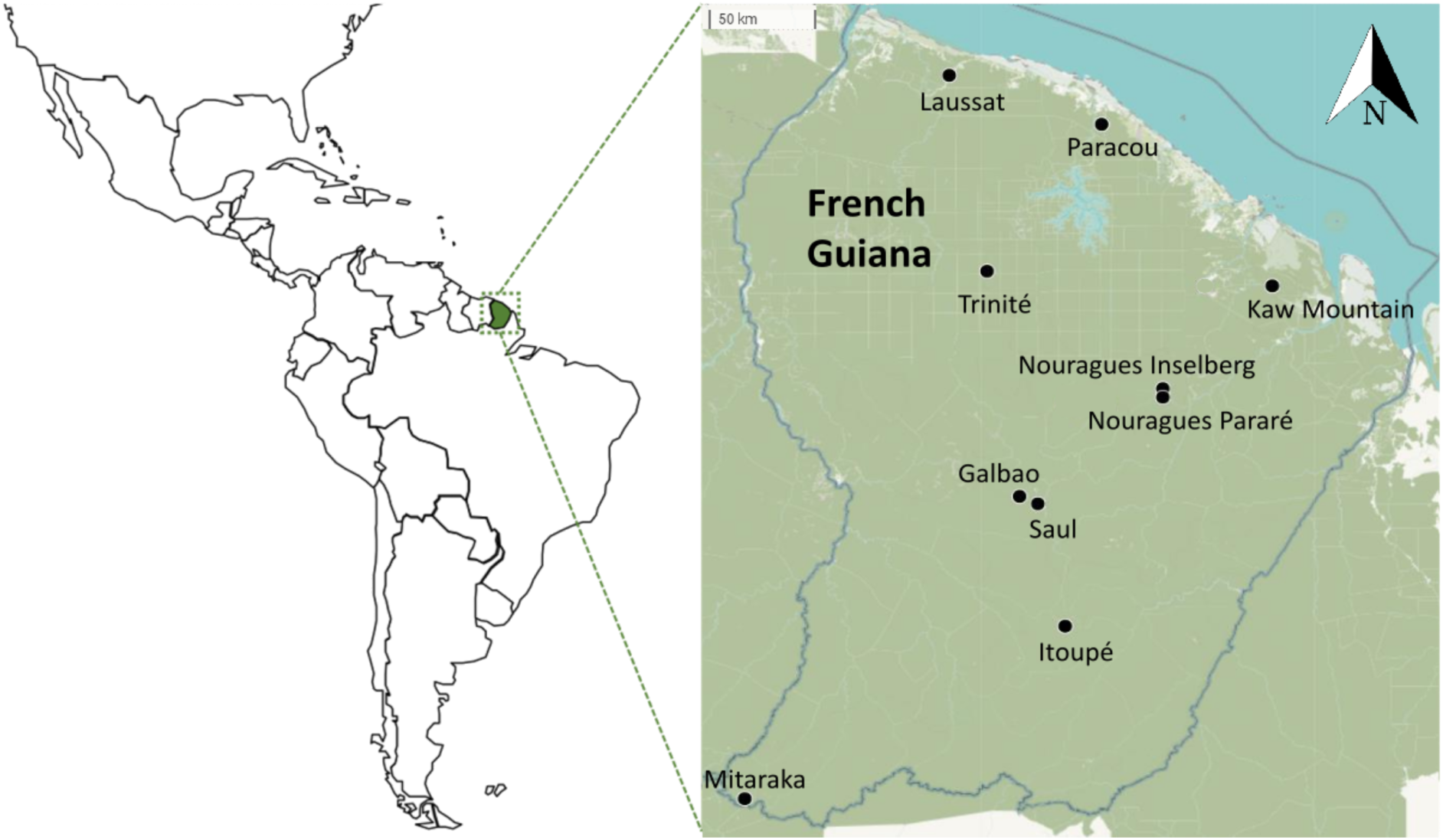
Location of the ten sampling localities in French Guiana.

In each locality, sampling was carried out in seven to twenty-five one-hectare plots, spaced at least 500 m apart, and representing replicates of the main habitat types of the local landscape (SI Table S1 and S2). Habitats included plateau forests (i.e. dense rainforest on deep, well-drained soils, sometimes with superficial laterites), slope forests (i.e. rainforest on slopes and deep soils), lowland forests (i.e. periodically flooded rainforest on hydromorphic soils) and plant associations specific to inselbergs and other landforms such as rocky savannas (i.e. herbaceous formations dominated by terrestrial bromeliads), and transition to hilltop forests (i.e. low canopy rainforest on shallow soils).

### Earthworm sampling

Sampling protocols have been described in Goulpeau, Hedde, Maggia, et al. (2024). Briefly, sampling was carried out on each one-hectare plot during the rainy season using a combination of three methods: 1) hand sorting of three blocks of soil of 25 × 25 cm^2^ area and 20 cm depth, spaced 20 m apart; 2) hand sorting of a 1 m^2^ surface to a minimum depth of 40 cm, selecting where possible a place with large earthworm surface casts; 3) active search in all available and accessible microhabitats (i.e. decomposing trunks, litter accumulations, streamside sediments, epiphytes, etc.) on the one-hectare plot for a fixed period of three hours.persons (e.g. one hour for three people). Specimens from all life stages (i.e. adults, juveniles and cocoons) were collected, and fixed in 100% ethanol.

### DNA barcoding and OTU delimitation

Earthworms collected with a given method in a given microhabitat and plot were first sorted according to external morphological features (i.e. clitellum characteristics when adult, size and pigmentation). For each morpho-group obtained, up to 5 individuals per sample were kept for DNA barcoding targeting the Cytochrome C Oxidase I gene (COI). The complete molecular biology protocols have been described in Goulpeau et al. (2022). Briefly, 77.2% of samples were Sanger processed according to standard protocols of the International Barcode of Life using a cocktail combining the M13 tail primer pairs LCO1490/HCO2198 (Folmer et al., 1994) and LepF1/LepR1 (Hebert et al., 2004). Failure tracking was done using the internal primers MLepR1/MLepF1 and LCO/HCO pairs (Hajibabaei et al., 2006). The remaining 22.8% sequences were obtained on MiSeq using LCO1490/HCO2198 primer pair (Folmer et al., 1994) combined with a tag on the 5’ side to allow further sample identification. Bioinformatic analysis was performed using OBITools (Boyer et al., 2016; www.grenoble.prabi.fr/trac/OBITools). Sequences were further grouped into molecular operational taxonomic units (OTUs) using the Assemble Species by Automatic Partitioning method (ASAP: Puillandre et al., 2021; https://bioinfo.mnhn.fr/abi/public/asap), which is the most recommended method for large numbers of earthworm DNA barcodes (Goulpeau et al., 2022). The validity of using these OTUs as surrogates for biological species is supported by morpho-anatomical characters obtained for the samples from two localities (Decaëns et al., 2016, 2024). All sequences are available in the BOLD dataset "Earthworms from French Guiana -2023 update" (DS-EWFG2023; doi.org/10.5883/DS-EWFG2023).

### Environmental parameters

Soil properties were described in a sub-sample of lowland, slope and plateau forest plots in Itoupé, Mitaraka, Saül and Trinité. In each of these plots, ten 0-30 cm deep soil cores were taken every 20 m along a transect passing through each plot (Vleminckx et al., 2019). The soil cores were combined and blended to form a composite sample from which a 500 g aliquot was taken, dried at 25 °C and sieved to 2 mm. Physico-chemical analyses were carried out at CIRAD laboratory, Montpellier France (https://us-analyses.cirad.fr) according to the protocols available on their website (Pansu & Gautheyrou, 2006). The variables measured were soil texture (mass proportion of clay, silt and sand), pH H_2_O, organic carbon (in mg.g^-1^ dry soil), total nitrogen (in mg.g^-1^ dry soil), C:N ratio and exchangeable phosphorus (Olsen method; in mg.kg^-1^ dry soil). Woody vegetation was described by calculating the abundance of trees grouped by family using data from (Vleminckx et al., 2019). Annual cumulative rainfall levels were obtained from the "POWER" database (https://power.larc.nasa.gov/data-access-viewer).

Elevation was recorded in the field using a GPS device, and the topographical position was then described by calculating the elevational deviation, defined as the standardised deviation from the mean altitude of the locality:

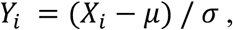

with *X_i_* the measured elevation of the plot, *μ* the average elevation of plots in the same locality, and *σ* the standard deviation from this average.

The patterns of variation in each of these parameters between localities and between habitat types are illustrated in SI Fig. S1.

### Dataset structure and diversity indices

We calculated species richness at a hierarchy of nested spatial extents: 1) alpha A is the richness at the community-scale, i.e. the number of OTUs found in a given one-hectare plot, 2) alpha B is the size of the habitat pool, i.e. the cumulated number of OTUs in a given habitat in a locality; 3) alpha C is the richness at the landscape scale, i.e. the number of OTUs found in a locality; 4) gamma is the regional richness, i.e. the total number of OTUs found in French Guiana (Table 1).

**Table 1.** Different levels of sampling hierarchy considered and definition of the diversity measures considered in the calculations of richness and beta diversity in the pairwise comparisons approaches.

| ***Spatial extent*** | ***Richness*** | ***Beta diversity*** |  |
| --- | --- | --- | --- |
| Community | **Alpha A:** number of OTUs/species in a given community | - |  |
| Habitat | **Alpha B:** number of OTUs/species in a given habitat and a given locality | **Beta A:** compositional dissimilarity among communities in a given habitat and a given locality |  |
| Locality (landscape) | **Alpha C:** number of OTUs/species in a given locality or landscape | **Beta B:** compositional dissimilarity among communities in different habitats in a given locality |  |
| Regional | **Gamma:** number of OTUs/species in the whole dataset | **Beta C:** compositional dissimilarity among communities from different localities or landscapes | **Beta D:** compositional dissimilarity among all communities |

Beta diversity was calculated between all possible pairs of communities with the Sørensen’s index of dissimilarity using the *’beta.pair’* function in the *’betapart’* package for R (Baselga et al., 2023; R Core Team, 2023):

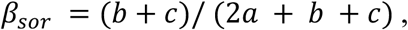

with *a* the number of species common to both communities, and *b* and *c* the number of species unique to the first and second community, respectively.

We calculated beta diversity for the following levels of data partitioning: beta A corresponds to intra-habitat beta diversity, i.e. the compositional dissimilarity between communities sampled in the same habitat and the same landscape, beta B corresponds to inter-habitat beta diversity, i.e. between communities sampled in different habitats within the same landscape, beta C corresponds to inter-landscape beta diversity, i.e. between communities sampled in different landscape. Ignoring the intermediate scales, beta D corresponds to regional beta diversity calculated for all possible community pairs in the dataset (Table 1). The mean values of beta diversity obtained for each of these partition levels were compared using a Welch t-test after first checking the normality and homoscedasticity of the data. This was done using the *’ggbetweenstats’* function in the *’ggstatsplot’* package for R (Patil, 2021).

### Alpha and beta diversity driving factors

In order to highlight the influence of biotic interactions on community-scale richness, we first adopted the popular approach of describing the relationship between local richness (alpha A) and species richness at higher spatial extents (alpha B or alpha C). A linear relationship between local and higher scale richness is typically interpreted as the result of unsaturated communities in which richness is essentially constrained by species dispersal capacity, whereas the absence of such a relationship is considered to be the result of competition which limits the number of species that can co-exist in saturated communities (Cornell, 1993; Cornell & Karlson, 1997). As this approach has been criticised (Hillebrand, 2005; Loreau, 2000), we have also used log-ratio models, which can highlight community saturation without any scale bias (Szava-Kovats et al., 2013). These models study the relationship between the dispersion of local richness and the log of richness at higher scales, which is negative in the case of saturated communities.

The links between community richness (alpha A) and environmental parameters were studied on the subset of thirty-six plots for which soil properties were available. We performed a PCA in order to select environmental variables not excessively correlated with each other. This resulted in removing from the analyses the percentage of clay and silt (that were highly correlated with each other and inversely correlated with sand content) and the total nitrogen content (that was correlated with both C content and C:N ratio). To account for the relative importance of selected variables as explanatory factors of alpha A, we used a generalised linear model (GLM: Dobson & Barnett, 2018; McCullagh & Nelder, 1989) using a Poisson distribution with the following formula:

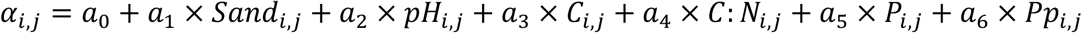

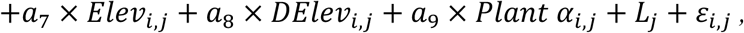

with *α_i,j_* the species richness of community *i*, which belongs to locality *j*. All factors indexed by *i,j* are measured in community *i*. *L_j_* is a random effect of the locality, distributed normally with mean 0 and standard deviation *σ_L_* and ε*_i,j_* is the individual error term, distributed normally with mean 0 and standard deviation *σ*. This was done using the *’glmer’* function in the *’lme4’* package for R (Bates et al., 2015).

We then used a generalised dissimilarity model (GDM: Ferrier et al., 2007) to highlight the relative importance of environmental dissimilarity and geographical distance as explanatory factors of beta diversity at the different levels of partition. We fitted these models by considering and then excluding geographical distance among communities. Environmental variables were the same as those used in the GLM, with the exception of vegetation, for which we used tree family composition, allowing us to consider plant beta diversity when calculating environmental dissimilarity. The model returns the percentage of deviance explained by geographical distance and environmental dissimilarity, as well as the weight of each variable or *’predictor importance’* and for each of them a p-value obtained using a permutation test (n = 100). These analyses were carried out using the *’gdm.varImp’* function in the *’gdm’* package for R (Ferrier et al., 2007), and the results were visualised using the *’corrplot’* package (Wei & Simko, 2021).

## Results

### Species richness

We were able to delimit 256 OTUs of earthworms across the 10 study localities. We observed an average richness of 7.0 OTUs at the community scale (alpha A), 15.9 OTUs at the habitat level (alpha B) and 33.2 OTUs at the landscape scale (alpha C) (Fig. 2A). Community-level richness ranged from one to 17 OTUs across the dataset, but 82% of assemblages at this scale contained fewer than 10 OTUs, and we found more than 15 OTUs in only one community (Fig. 2B). We found a significant linear relationship between alpha A and the size of habitat species pool (alpha B) (Fig. 2B). Furthermore, we observed a negative relationship (Fig. 2C) between the dispersion of alpha A and both alpha B and alpha C based on a log-ratio regression model (Szava-Kovats et al., 2013).

**Figure 2.**
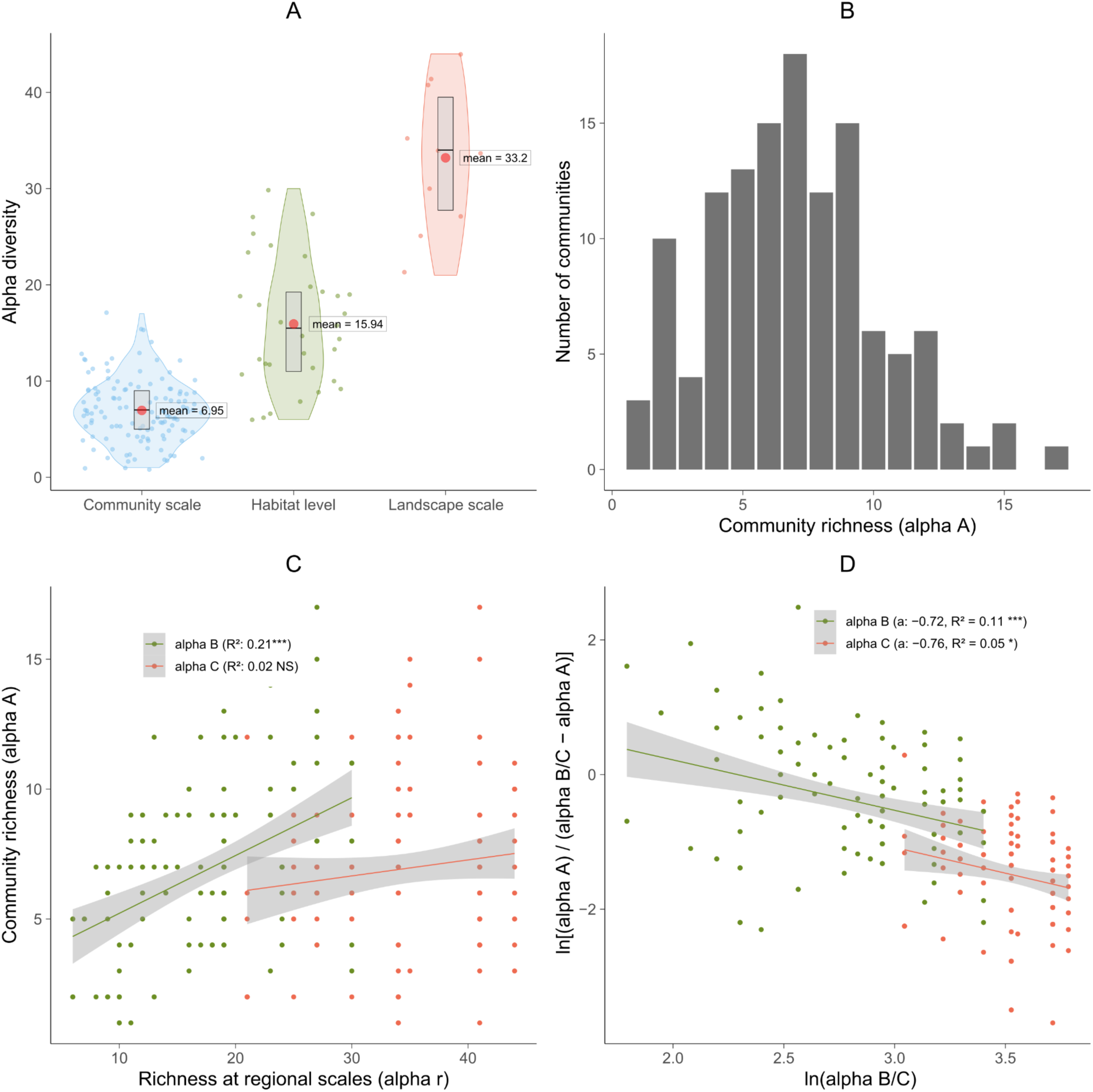
Earthworm community richness in rainforests of French Guiana: A) Species richness at the community scale (alpha A), habitat level (alpha B) and landscape scale (alpha C); B) Number of communities per richness level; C) Linear models of the relationship between community-scale richness (alpha A) and richness at higher spatial extents, i.e. alpha B or alpha C ; D) Log-ratio models of the relationship between the dispersion of community-scale richness and the log of species richness at higher spatial extents (Szava-Kovats et al., 2013). In C and D, lines represent the fitted models, shaded areas are the 95% confidence interval, a = regression coefficient. Model significance codes: 0 ’***’ 0.001 ’**’ 0.01 ’*’ 0.05 ’.’ 0.01.

GLM analysis revealed that soil texture (% sand) was the only environmental factor that significantly explained community-level alpha diversity, with alpha A decreasing with the sand content in the soil (Fig. 3; SI, Table S3).

**Figure 3.**
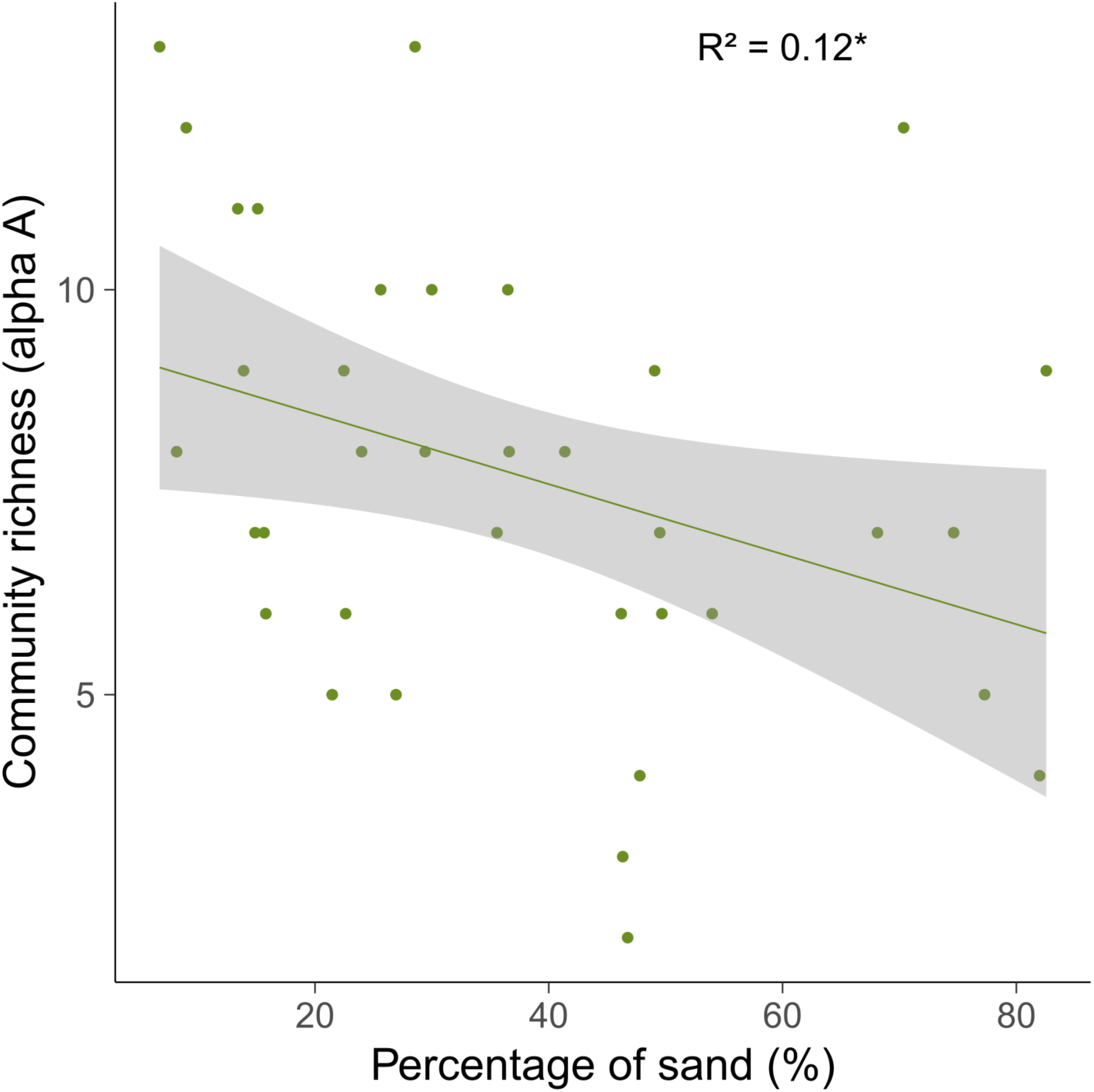
Relationship between soil texture and community-scale richness. Other environmental factors tested in the generalised linear model were not significant and are not shown graphically. The line represents the fitted linear model; the shaded area is the 95% confidence interval; p < 0.1.

### Beta diversity

Beta diversity between pairs of communities averaged 0.89 for the whole dataset (beta D), and varied significantly between partition levels (Fig. 4). It was the lowest between communities sampled in the same habitats and the same landscape (beta A), increased significantly for communities sampled in different habitats (beta B), and it was the highest when the communities being compared belonged to different localities (beta C).

**Figure 4.**
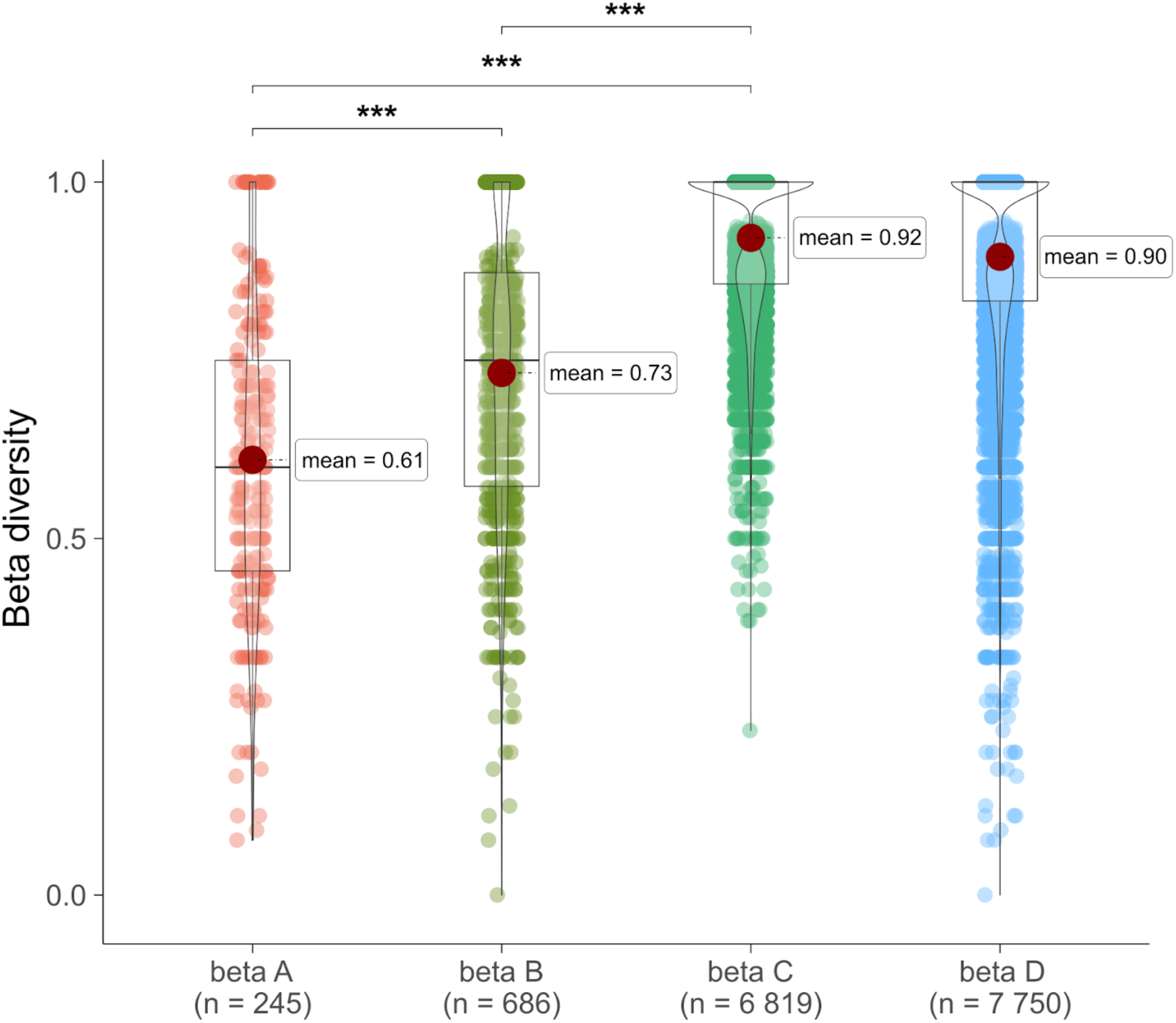
Sørensen’s beta diversity between pairs of earthworm communities partitioned into different levels of spatial extent: beta A is the beta diversity between communities within the same habitat of the same landscape, beta B is the beta diversity between communities of different habitats of the same landscape, beta C is the beta diversity between communities located in different landscapes, beta D is the beta diversity between all possible pairs of communities within the dataset. Pairwise test Games-Howell significance codes : 0 ’***’ 0.001 ’**’ 0.01 ’*’ 0.05 ’.’ 0.1.

GDMs allow to highlight the relative importance of geographical distance and environmental dissimilarity in explaining beta diversity between pairs of communities, and to identify the driving factors of this beta diversity across the hierarchy of partition levels (Fig. 5). At the habitat-level (beta A), the models explained 42.5% and 30.6% of the deviance (when geographical position was taken into account or not, respectively), and geographical distance was the only significant predictor of beta diversity. When comparing communities sampled in different habitats of the same landscape (beta B), the models explained 38% and 37.2% of the deviance in beta diversity, which was essentially explained by altitudinal deviation and soil assimilable phosphorus content. On a regional scale, the two models explained only 13.4% of the deviance when pairs of communities located in different localities were taken into account (beta C), and no individual factor was significant. However, when all possible pairs of communities were considered (beta D), the models explained 31.1% and 23.6% of the deviance, and distance appeared to be the main predictor of beta diversity. When geographical position was removed from the model, mean annual precipitation and elevation appeared as significant predictors, suggesting that these two factors exhibit strong spatial autocorrelation at this scale.

**Figure 5.**
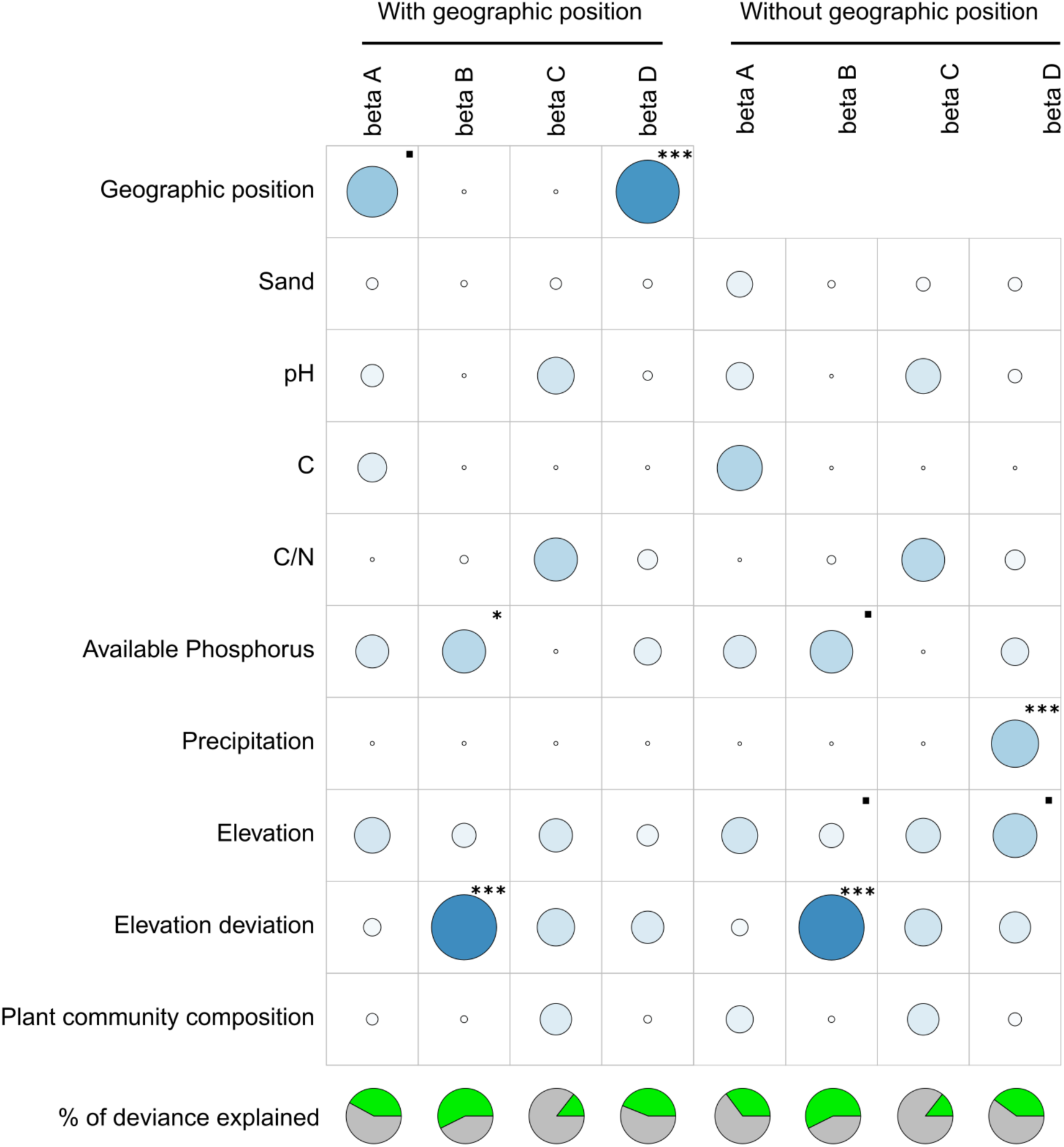
Importance of geographical position and environmental factors as explanatory factors of beta diversity between pairs of communities at different partition levels (as explained in Table 3). Generalised dissimilarity models did or did not consider distance between communities as a spatial factor (with or without geographic position, respectively). Surface areas of circles are proportional to the relative contribution of each factor to this model. Pie charts show in green, for each level of beta diversity, the percentage of deviance explained by the model; sand= % sand in soil texture; C= soil C content. GDM significance codes : 0 ’***’’**’ 0.01 ’*’ 0.05 ’.’ 0.1.

## Discussion

### Competition and soil properties explain community-scale alpha diversity

On average, we observed only seven OTUs per community, and rarely more than 10 OTUs in a given community. This is in line with other studies which show that earthworm communities rarely comprise more than ten species, with little or no effect of latitude and of the size of regional species pools (Lavelle & Spain, 2001; Wardle, 2002). They are also in line with the findings of Phillips et al. (2019), who highlighted low values of local richness for earthworm communities in most tropical regions. Such a limitation of community-scale alpha diversity suggests that competition may constrain the number of species co-existing locally in a context of resource limitation, this independently of the level of landscape diversity (Alroy, 2018; Hillebrand, 2005; Loreau, 2000).

This interpretation seems to be contradicted by the linear relationship observed between community and habitat or landscape richness, which is classically interpreted as reflecting unsaturated communities in which species dispersal capacity is the main factor explaining local diversity (Cornell, 1993; Hillebrand, 2005; Loreau, 2000). However, the use of such regressions to detect community saturation has been criticised due to the existence of a scale bias (Hillebrand, 2005; Loreau, 2000). Szava-Kovats et al. (2013) proposed that the use of log-ratio models could overcome this bias and reveal more efficiently saturated communities.

When fitting these models to our data, we found a significant slope between alpha A and alpha B, revealing communities that are at least partially saturated. This result is in line with other studies that have already highlighted earthworm community patterns attributable to competition effect, such as character displacement (Fragoso & Rojas, 1997), competitive displacement (Callaham et al., 2003; Winsome et al., 2006), spatial structuring of species assemblages (Decaëns & Rossi, 2001; Jimenez et al., 2006), lower co-occurrence than expected by chance (Decaëns et al., 2008) and the aforementioned lack of relationship between local and regional diversity (Lavelle & Spain, 2001; Wardle, 2002).

Our results also show that community richness is at least partly determined by the local environment. In particular, we observed a negative correlation between alpha A and soil sand content. These results are broadly in line with recent meta-analyses that have identified soil texture and climate as important drivers of earthworm diversity on a global scale (Phillips et al., 2019). However, we do not find the overriding importance of annual rainfall highlighted by Phillips et al. (2019). This difference can be explained by the more restricted spatial extent of our study, which covers a limited range of environmental conditions compared to global studies: climate in French Guiana is supposed to vary relatively little compared with the variations that can be observed by comparing different localities on a more global scale. Similarly, we did not observe any link with tree composition, even though it has been shown that vegetation diversity and composition can be a key factor determining temperate earthworm diversity, mainly because of its link with the trophic resources available (Cesarz et al., 2007; Ganault et al., 2021). Here again, this difference can be explained by the fact that vegetation diversity in French Guiana is high regardless of the study site, buffering the effects of plant composition on the quality of the resource for soil fauna.

Overall, these results support our second hypothesis that community-level richness is explained both by local environment and interspecific interactions. The levelling off of community richness, and the relationship between local and regional richness, suggest firstly that communities are at least partially saturated. However, the local environment may also limit community richness, particularly in the sandier soils, which offer the least favourable conditions for earthworms in terms of water resources.

### Distance and environment as drivers of beta diversity among pairs of communities

The fact that the composition of species assemblages is more similar between communities in the same habitat (beta A), and that Sørensen’s dissimilarity increases consistently with environmental distance (beta B) and with geographical distance between communities (beta C), suggests the existence of both local habitat signal and regional geographical turnover as already proposed by Maggia et al. (2021). At each of these partition levels, the GDMs allow us to further identify the factors responsible for these patterns in compositional dissimilarity between communities.

On a regional scale, Sørensen’s dissimilarity calculated over our entire dataset (beta D) is consistently high, and essentially explained by the geographical distance between communities. Furthermore, when removing the spatial effect from the model, the effect of altitude and rainfall become significant. When only pairs of communities from different landscapes are taken into account (beta C), however, this effect disappears, probably because the range of variation in geographical distances is too small in this level of partition. The relationship between beta D and geographical distance is a direct result of the strong compositional dissimilarity between the species pool composition of distant landscapes that has been highlighted by Goulpeau, Hedde, Ganault, et al. (2024). Given that community composition represents a subsample of the species pool within a specific landscape, high compositional dissimilarity between landscapes species pools inevitably results in high beta diversity between pairs of distant communities. The fact that precipitation and altitude appear as explanatory factors when the spatial effect is removed from the model indicates that these factors are spatially autocorrelated, and that they both explain beta diversity at this scale. French Guiana is characterised by a regional rainfall gradient that is partly correlated with the relief, with the south generally more elevated and less rainy than the north (Barret, 2001). In addition to the isolation of populations already mentioned, this regional climatic variability may have favoured vicariance events by locally modifying selective pressures, thereby accentuating the differentiation of species pools at this regional scale. These results are broadly in line with the multi-taxa study by (Dambros et al., 2020), who conclude that the composition of communities in Amazonia reflects both the limitation of dispersal (isolation by distance and by major rivers) and the adaptation of assemblages to local environmental conditions.

The significant difference between average beta diversity between communities of different habitats (beta B) and that between communities of the same habitats (beta A) illustrates the effect of environmental filters on community assembly. The environment selects from the landscape pool the species presenting the necessary adaptations to settle successfully in a local habitat, leading locally to communities that are more similar in a given habitat than between different habitats. The GDMs show that beta diversity between habitats is associated with topography (i.e. elevational deviation) as well as with differences in total phosphorus availability. Phosphorus is a key limiting element in tropical rainforests, whose availability in soil and litter regulates important ecosystem processes such as primary productivity, litter production and decomposition, and nutrient cycling in general (Cleveland et al., 2011). Differences in available phosphorus in soils from one habitat to another can lead to heterogeneity in vegetation composition (Vleminckx et al., 2019; Wright et al., 2011), with repercussions on the diversity and quality of trophic resources for soil organisms (Fichaux et al., 2021; McGlynn et al., 2007; Vleminckx et al., 2019). The fact that vegetation composition did not emerge in our results as a factor controlling beta diversity seems to contradict this interpretation, but it is possible that the way in which we took vegetation into account in our models (i.e. family composition) does not fully account for its effect on earthworm resources. Future studies could consider the effect of vegetation structure and/or litter traits to better explore this mechanism.

When intra-habitat beta diversity is considered (beta A), the GDMs highlight distance between communities as the only significant explanatory factor, unlike all the environmental factors considered in the model. This suggests that community assembly within a given local habitat is essentially governed by neutralism (Chave, 2004). This has previously been demonstrated in tropical plant communities, where dispersal limitation alone predicts a substantial fraction of beta-diversity (Condit et al., 2002; Hardy & Sonké, 2004). In earthworms, it has been shown that communities are spatially structured into opposing assemblages that may result from the spatial segregation of competing species and/or the response of species to patchiness of soil properties (Decaëns & Rossi, 2001; Jiménez et al., 2012). On the other hand, Rossi et al. (1997) also demonstrated that the spatial structuring of earthworm populations can result from neutral mechanisms involving solely demographic dynamics and dispersal limitation. However, these studies focused on assemblage structuring within communities, whereas we go further by demonstrating that neutral mechanisms explain the compositional dissimilarity of communities in the same habitat on a landscape-scale.

## Conclusion

Our study provides new insights into the factors controlling earthworm diversity in tropical rainforests. In accordance with our first hypothesis, we found that alpha diversity at the community-level is explained both by local environment (soil texture) and by competition, which limits the number of species that can coexist in saturated communities. Our results also partly support our second hypothesis. Indeed, we highlight that habitat filtering linked with topography and soil phosphorus availability explains the compositional dissimilarity between communities from one habitat to the other. We also found that beta diversity among all possible pairs of communities is essentially explained by distance and climate, suggesting that the spatial distribution of species diversity at this regional scale is the result of biogeographic processes involving dispersal limitation and local adaptations to environmental conditions. The beta diversity between communities in the same habitat, although lower than that observed for the other partition levels in our study, is nevertheless higher than we had initially thought. It is explained exclusively by the distance between communities, which suggests a degree of neutralism in the community assembly process. To go a step further in analysing the assembly rules of tropical earthworm communities, it will be interesting in future studies to consider functional and phylogenetic diversity in an approach similar to ours. Combining trait-based and phylogenetic approaches of biodiversity would indeed represent a powerful strategy to disentangle the ecological and evolutionary drivers of community structure (Emerson & Gillespie, 2008), and to improve our understanding of large-scale variations in species diversity distribution (Lamanna et al., 2014; Mouquet et al., 2012; Violle et al., 2014) and its vulnerability to on-going global changes (Lavergne et al., 2010; Mouillot et al., 2013).

## Acknowledgements

Part of the dataset used in this study was acquired as part of the DIADEMA (Dissecting Amazonian Diversity by Improving a Multiple Taxonomic Group Approach), DIAMOND (Dissecting and Monitoring Amazonian Diversity) and Wormbank (DNA barcoding earthworms in biodiversity hot spots of French Guiana) projects funded by "Investissement d’Avenir" grants managed by the Agence Nationale de la Recherche (CEBA: ANR-10-LABX-25-01; TULIP: ANR-10-LABX-41). Sampling in Mitaraka was carried out as part of the "Our Planet Reviewed" French Guiana 2015 expedition (Touroult et al., 2018) organised by the Muséum national d’Histoire naturelle (MNHN, Paris) and Pro-Natura international in collaboration with the Amazonian Park of French Guiana, and financed by the European Fund for Regional Development (FEDER), the Regional Council of French Guiana, the General Council of French Guiana, the Direction de l’Environnement, de l’Aménagement et du Logement and by the Ministère de l’Éducation nationale, de l’Enseignement supérieur et de la Recherche. At the Réserve naturelle des Nouragues, the project was supported by two Centre National de la Recherche Scientifique (CNRS) Nouragues grants in 2010 and 2011. At the Réserve naturelle de la Trinité, part of the funding was provided by the nature reserve. The authors would like to thank the Parc Amazonien de Guyane (http://www.parcamazonien-guyane.fr), the Réserve naturelle de la Trinité (http://www.reserve-trinite.fr/) and the Réserve naturelle des Nouragues (http://www.nouragues.fr/) for authorising access and collecting. Sample DNA barcoding received funding from the Canadian Centre for DNA Barcoding (CFREF-2015-00004), as well as from the CNRS through the BC-Wormbank project (APEGE 2013 call).

## Author contributions

AG analysed the data and led the writing of the manuscript. MH and TD supervised the study. TD and EL led field sampling and sample processing. All authors contributed to the writing of the manuscript.

## Data availability statement

Specimens list, metadata, COI sequences and GenBank accession numbers are available in the BOLD dataset “Earthworms from French Guiana -2023 update” (DS-EWFG2023; doi.org/10.5883/DS-EWFG2023). Specimens list, community table and environmental data have been published in Goulpeau, Hedde, Maggia, et al. (2024), and are deposited on the Zenodo repository (doi.org/10.5281/zenodo.10908657).

## Supporting information

Appendix S1.

Appendix S2.

## Dissecting earthworm diversity in tropical rainforests

Arnaud Goulpeau, Mickaël Hedde, Pierre Ganault, Emmanuel Lapied, Marie-Eugënie Maggia, Eric Marcon, Thibaud Decaëns

## Appendix S1. Material and method supporting information

**Table S1.** List of sampled landscapes, average geographical position, sampling periods and number of communities sampled in each.

| Landscapes | Latitude | Longitude | Sampling data | Number of communities |
| --- | --- | --- | --- | --- |
| Laussat | 5.48117 | -53.5762 | 02/2014 - 02/2018 - 12/2019 | 7 |
| Paracou | 5.25833 | -52.9346 | 06/2013 | 10 |
| Trinité | 4.61871 | -53.4081 | 04/2016 | 10 |
| Kaw | 4.54064 | -52.2152 | 06/2013 - 04/2016 | 9 |
| Inselberg | 4.08133 | -52.6731 | 01/2011 - 06/2011 - 02/2018 | 25 |
| Pararé | 4.02832 | -52.6778 | 06/2011 - 07/2011 - 02/2018 | 12 |
| Galbao | 3.60403 | -53.268 | 01/2016 - 01/2019 | 11 |
| Saül | 3.5587 | -53.2222 | 10/2013 - 02/2014 | 12 |
| Itoupé | 3.02696 | -53.079 | 01/2016 | 11 |
| Mitaraka | 2.243 | -54.465 | 03/2015 | 8 |

**Table S2.**
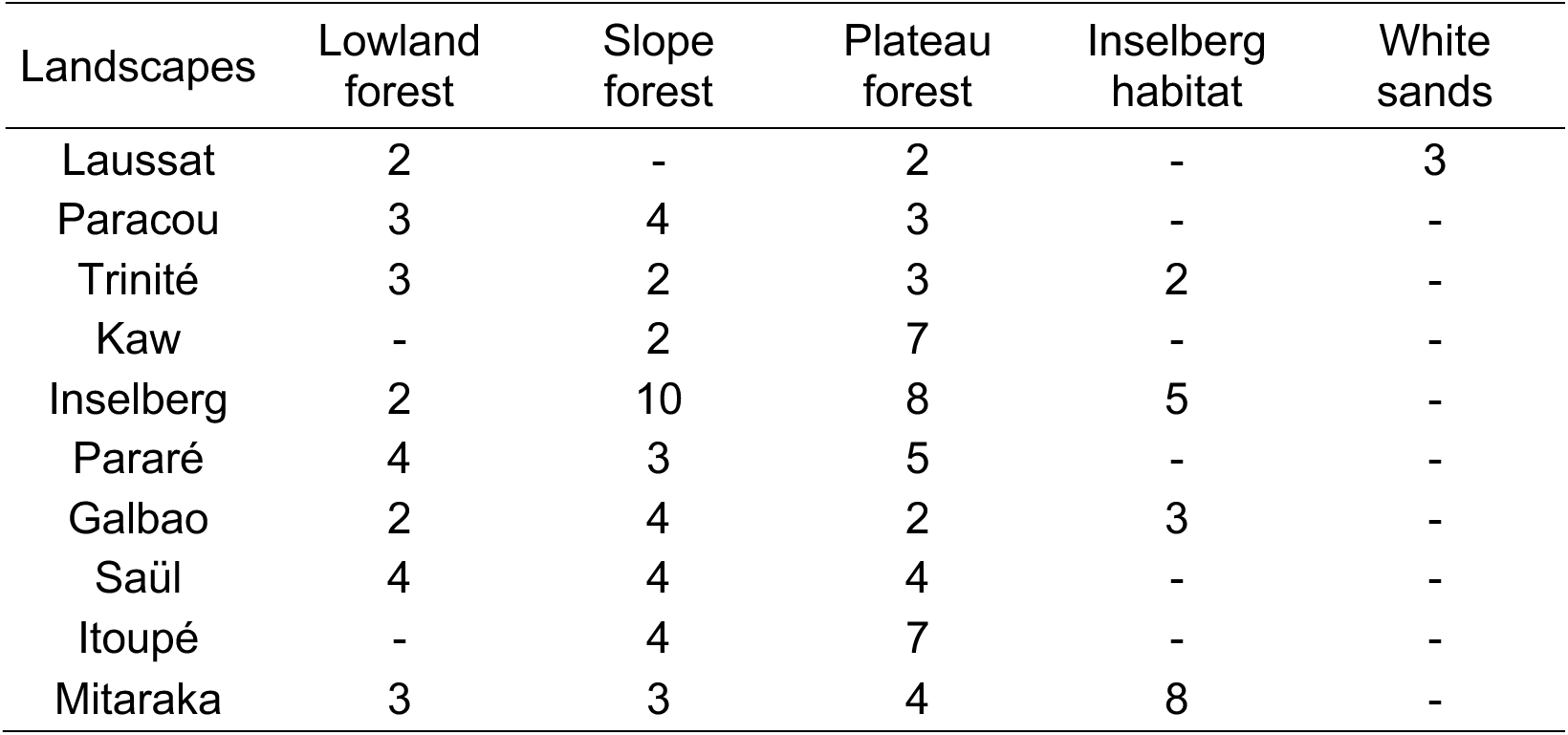
Number of communities sampled per type of habitat in each landscape.

| Landscapes | Lowland forest | Slope forest | Plateau forest | Inselberg habitat | White sands |
| --- | --- | --- | --- | --- | --- |
| Laussat | 2 | - | 2 | - | 3 |
| Paracou | 3 | 4 | 3 | - | - |
| Trinité | 3 | 2 | 3 | 2 | - |
| Kaw | - | 2 | 7 | - | - |
| Inselberg | 2 | 10 | 8 | 5 | - |
| Pararé | 4 | 3 | 5 | - | - |
| Galbao | 2 | 4 | 2 | 3 | - |
| Saül | 4 | 4 | 4 | - | - |
| Itoupé | - | 4 | 7 | - | - |
| Mitaraka | 3 | 3 | 4 | 8 | - |

**Figure S1.**
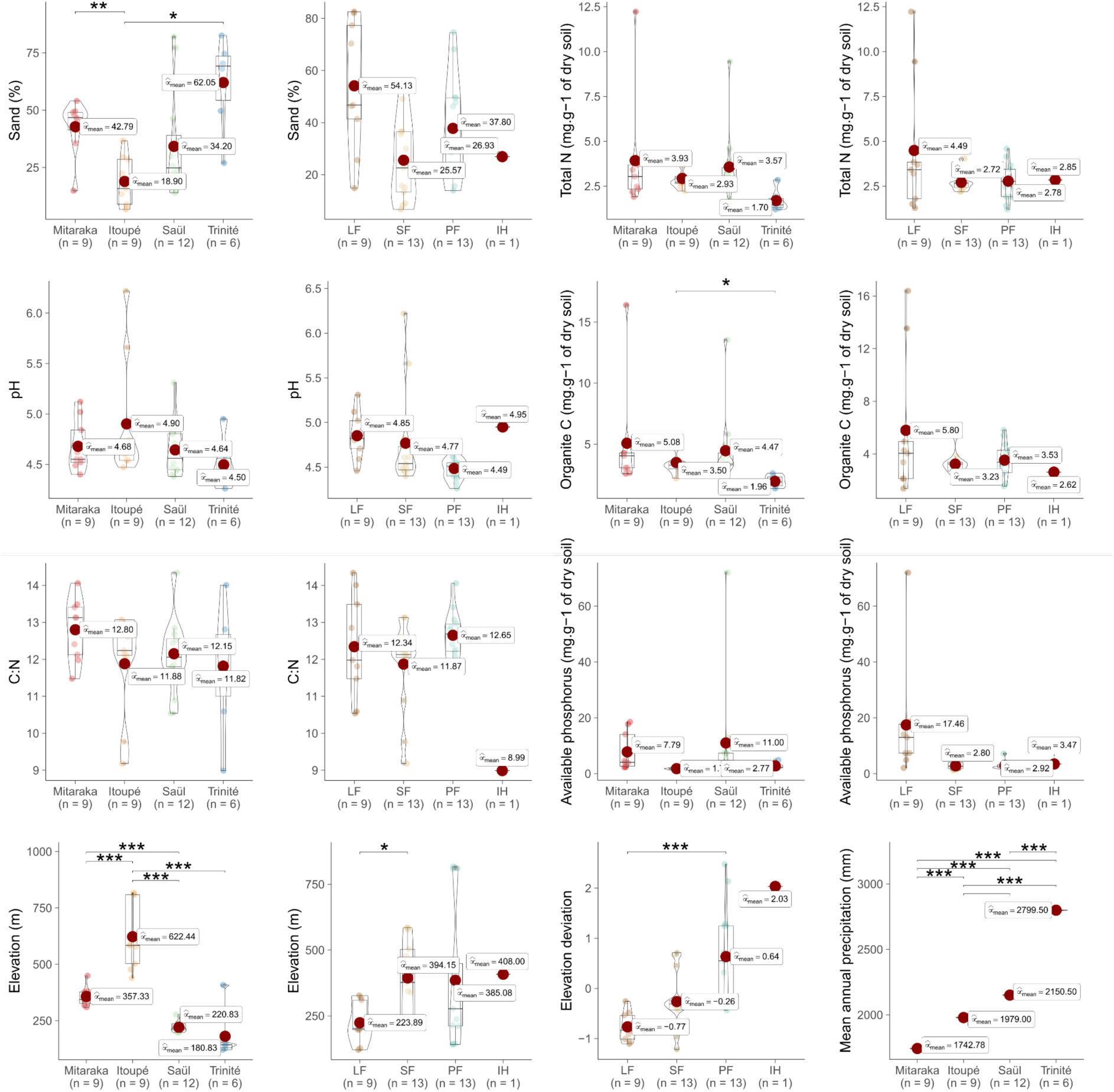
Box-plots showing patterns of variation in average environmental factors between landscapes and habitat types. LF= lowland forest; SP= slope forest; PF= plateau forest; IH= inselberg habitats. Pairwise test Games-Howell significance codes : 0 ’***’ 0.001 ’**’ 0.01 ’*’0.05 ’.’ 0.1. Dataset available on Zenodo (doi.org/10.5281/zenodo.8410359).

**Figure S2.**
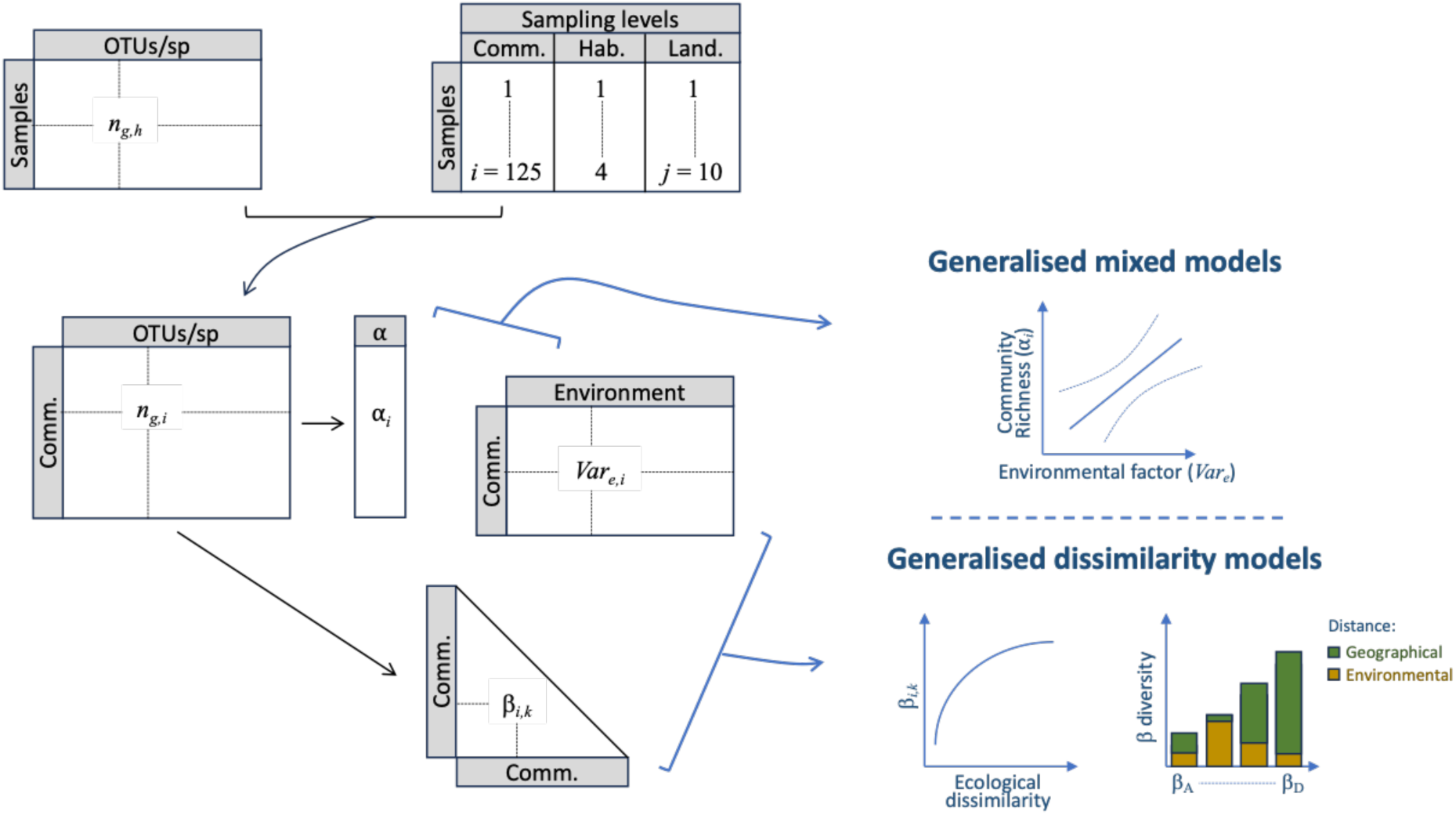
Diagram summarising dataset structure and analysis strategy. The data tables are indicated in grey, and analyses are indicated in blue. In datasets, *h* is the number of samples, *i* is the number of sampled communities, *j* is the number of landscapes, *e* is the number of environmental variables, *g* is the number of operational taxonomic units of species level (OTUs), *n_gh_* is the number of specimens of the OTU *g* collected in the sample *h*, *_ngi_* is the number of specimens of the OTU *g* collected in the community *i*, **α*_i_* is the number of OTUs in the community *i* (i.e. community-scale richness), *β_ik_* is the Sørensen’s beta diversity among communities *i* and community *k*, *Var_ei_* is the value taken by the environmental variable *e* in the community *i*; sampling hierarchy levels are: communities (Comm.), habitats (Hab.) and landscapes (Land.). Generalised mixed models are used to identify the environmental drivers of community-scale **α** diversity. Generalised dissimilarity models are used to highlight the relationships between *β* diversity among pairs of communities and ecological dissimilarity, which is partitioned into the effects of environmental and geographical distances at the different levels of sampling hierarchy (*β_A_* to *β_D_*).

## Appendix S2. Result supporting information

**Table S3.** Generalised linear model results: coefficients and significance of the environmental variables as explanatory factors of earthworm species richness at the community level (alpha 2). Sand = proportion of sand in soil texture; pH = soil pH (H_2_O); C = total carbon content of soil; C:N = carbon to nitrogen ratio in soil.

|  | Estimate | Std. Error | z value | Pr(>\|z\|) |
| --- | --- | --- | --- | --- |
| (Intercept) | -0,3876 | 2,4833 | -0,1561 | 0,8760 |
| Sand | -0,0089 | 0,0042 | -2,1289 | 0,0333* |
| pH | 0,0583 | 0,2790 | 0,2090 | 0,8344 |
| C | -0,0346 | 0,0423 | -0,8166 | 0,4142 |
| C/N | 0,1228 | 0,0916 | 1,3406 | 0,1800 |
| Available Phosphorus | 0,0008 | 0,0099 | 0,0798 | 0,9364 |
| Precipitation | 0,0004 | 0,0003 | 1,7333 | 0,0830 |
| Elevation | 0,0004 | 0,0005 | 0,9114 | 0,3621 |
| Elevation deviation | -0,0667 | 0,0772 | -0,8636 | 0,3878 |
| Plant alpha diversity | 0,0013 | 0,0126 | 0,1033 | 0,9177 |

